# Targeting ON-bipolar cells by AAV gene therapy stably reverses *LRIT3*-congenital stationary night blindness

**DOI:** 10.1101/2021.10.04.462588

**Authors:** Keiko Miyadera, Evelyn Santana, Karolina Roszak, Sommer Iffrig, Meike Visel, Simone Iwabe, Ryan F. Boyd, Joshua T. Bartoe, Yu Sato, Alexa Gray, Ana Ripolles Garcia, Valerie Dufour, Leah C. Byrne, John G. Flannery, William A. Beltran, Gustavo D. Aguirre

## Abstract

AAV gene therapies aimed at curing inherited retinal diseases to date have typically focused on photoreceptors and retinal pigmented epithelia within the relatively accessible outer retina. However, therapeutic targeting in diseases such as congenital stationary night blindness (CSNB) that involve defects in ON-bipolar cells (ON-BCs) within the mid-retina has been challenged by the relative inaccessibility of the target cell in intact retinas, the limited transduction efficiency of these cells by existing AAV serotypes, poor availability of established ON-BC-specific promoters, and absence of appropriate patient-relevant large animal models. Here, we demonstrate safe and effective ON-BC targeting by AAV gene therapy in a recently characterized naturally-occurring canine model of CSNB, *LRIT3*-CSNB. To effectively target ON-BCs, new AAV capsid variants with ON-BC tropism and ON-BC specific modified GRM6 promoters were adopted to ensure cell-specific transgene expression. Notably, subretinal injection of one vector, AAV^K9#4^-*sh*GRM6-c*LRIT3*-WPRE, significantly recovered rod-derived b-wave in all treated eyes (6/6) of adult dogs injected at 1-3 years of age. The robust therapeutic effect was evident 7 weeks post-injection and was sustained for at least 1 year in all treated eyes. Scotopic vision was significantly improved in treated eyes based on visually-guided obstacle course navigation. Restoration of LRIT3 signals was confirmed by immunohistochemistry. Thus, we report on the first ON-BC functional rescue in a large animal model using a novel AAV capsid variant and modified promoter construct optimized for ON-BC specificity, thereby establishing both proof-of-concept and a novel translational platform for treatment of CSNB in patients with defects in photoreceptor-to-bipolar signaling.

**Significance:** Canine models of inherited retinal diseases have informed and advanced AAV gene therapies targeting specific cells, including photoreceptors in the outer retina, for treatment of blinding diseases in human patients. However, therapeutic targeting of diseases such as congenital stationary night blindness (CSNB) that exhibit defects in ON-bipolar cells (ON-BCs) of the mid-retina remains under-developed. Using a new *LRIT3* mutant canine model of CSNB exhibiting ON-BC dysfunction, we tested the ability of a cell-specific AAV capsid and promotor to specifically target ON-BCs for gene delivery. Notably, subretinal injection of AAV-*LRIT3* vector demonstrated safety and efficacy with robust and stable rescue of ERG signals and night vision up to 1 year, paving the way for clinical trials in CNSB patients.

## Introduction

The vision process begins with photon capture by outer segments of rods and cones, photoreceptors within the outer retina. Subsequent release of neurotransmitters by photoreceptor synaptic terminals – rod spherules and cone pedicles – transfers this information to bipolar cell dendritic processes in the outer plexiform layer. Notably, distinct cell classes and signaling pathways allow for day and night vision. Specifically, night vision requires signaling from rod photoreceptors to ON-bipolar cells (ON-BCs), and activation of the mGluR6 signaling cascade at the dendritic tips of ON-BCs. In the dark, glutamate is released from photoreceptor terminals, binds to the mGluR6 receptor, and activates the alpha subunit of a G-protein, Goα. Dissociation of G-protein subunits culminates in the closure of TRPM1 channels, the key regulator of the mGluR6 signaling cascade^1^. Several molecules, including LRIT3 and NYX, are critical in localizing TRPM1 to the ON-BC dendritic tip membrane^2^. Inherited defects in the pathway, from photon capture, phototransduction, neurotransmitter release and mGluR6 signaling cause non-progressive vision impairing disorders, termed congenital stationary night blindness (CSNB), that are characterized by the absence of night vision from birth.

The genetically heterogeneous causation of CSNB^3^ underlies the spectrum of clinical presentations, broadly classified using electroretinography (ERG) as Riggs^4^ and Schubert-Bornschein^5^ forms of CSNBs. The Riggs type is caused by defects in the photoreceptors and is associated to date with four genes, *RHO, GNAT1, PDE6B*, and *SLC24A1*, in decreasing order of prevalence^3^. The Schubert-Bornschein type results from erroneous photoreceptor-to-bipolar cell signaling and is further subdivided into incomplete and complete CSNB^6^. Incomplete CSNB is associated with partial blockage of photoreceptor-to-bipolar cell signaling that results in some residual rod function. Defects in three genes, *CACNA1F, CABP4*, and *CACNA2D4*, all expressed pre-synaptically in the photoreceptor terminals are associated with incomplete CSNB^3^. Complete CSNB (cCSNB) is the most common form and patients exhibit characteristic ERG features including extinguished scotopic b-wave due to ON-BC dysfunction. Defects in five genes, *NYX*^7,8^, *TRPM1*^9-11^, *GRM6*^12,13^, *GPR179*^14,15^, and *LRIT3*^16^, all historically thought to localize post-synaptically in the ON-BC dendritic tips, are associated with cCSNB.

A series of cCSNB mouse models with ON-BC defects are categorized as ‘*nob’* (no b-wave) mice^3^. Consistent with the non-progressive nature of CSNB, *Lrit3* deficient mice (*Lrit3*^-/-^, *Lrit3*^nob6^) have an intact outer nuclear layer (ONL)^17^, and the observed cone-specific disorganization of synaptic contacts support a role for LRIT3 in the transsynaptic communication between cones and ON-BCs^18^. As with other nob mouse models, *Lrit3*-deficient *nob6* mice only partially recapitulate the phenotype of *LRIT3*-cCNSB human patients, due to unexpected loss of photopic b-waves in addition to expected loss of scotopic b-waves (**Table S1**)^16,17^.

We have previously characterized a naturally-occurring canine model of CSNB with striking night blindness and normal day vision. Full-field ERGs showed a predominant loss of scotopic b-waves, consistent with Schubert-Bornschein type cCSNB^19^. A subsequent genome-wide association study and whole-genome sequencing identified an exonic 1bp insertion in *LRIT3*^20^, the leucine-rich repeat Immunoglobulin-like transmembrane domain 3 gene associated with cCSNB in both patients^16^ and the murine model^17^. The canine LRIT3 is predicted to have a signal peptide, two LRR (Leucine-rich repeats), two LRR type domains, and one each of LRRCT (LRR C-terminal), IGc2, FN3, and transmembrane domains. The canine *LRIT3* disease variant generated a premature stop codon, giving rise to a truncated LRIT3 predicted to lack IGc2, FN3, and C-terminal transmembrane domains^20^. *In vitro* studies showed subcellular expression and distribution of the truncated LRIT3 was comparable to wild type (WT) LRIT3, indicating although non-functional it is unlikely to be toxic. Moreover, immunohistochemistry (IHC) analysis showed punctate LRIT3 labeling in putative ON-BC dendritic terminals co-labeled with Goα in WT, but markedly decreased levels in the mutants^20^, suggesting *LRIT3* augmentation might be an effective approach for therapeutic intervention. To target ON-BCs more effectively in mature retina, we screened a small pool of novel AAV capsid variants for ON-BC tropism in adult dogs. Herein we report the first ON-BC functional rescue in a large animal model using a novel AAV variant and a modified promoter, both designed for ON-BC specificity. We establish proof-of-concept and provide a translational platform for treatment of CSNB driven by defects in photoreceptor-to-bipolar signaling.

## Results

### Validation of new AAV capsid variants

Two AAV capsid variants, AAV^K9#4^ and AAV^K9#12^ were selected following a screen for ON-BC targeting in dogs (**Fig. S1, S2**) and tested in a non-human primate (NHP) using a long version of modified mGluR6 (*GRM6*) promoter (*lg*GRM6)^21^. The AAV^K9#4^ and AAV^K9#12^ capsids were each selected based on their superior capacity for ON-BC specific expression when injected via subretinal and intravitreal routes, respectively. *In vivo* cSLO imaging in WT dogs injected with AAV^K9#4^-*lg*GRM6-*sfGFP* subretinally in one eye and AAV^K9#12^-*lg*GRM6-*sfGFP* intravitreally in the contralateral eye (3×10^13^ vg/mL), showed strong sfGFP fluorescent signal in the subretinally-treated bleb area and in the central to mid-peripheral retina of the intravitreally-injected eye (**Fig. 1A**). Further, IHC confirmed tropism of both vectors to ON-BCs (**Fig. 1B, C**). Quantification of Goα-positive cells showed that intravitreal AAV^K9#12^ transduced 34-86% (central), 0-45% (mid-periphery), and 0-3% (periphery) of ON-BCs (**Fig. 1C, D**). Transduction efficiency with AAV^K9#4^ in the subretinal bleb area was more homogeneous, ranging from 57% (periphery) to 80% (center) (**Fig. 1B, D**).

**Figure 1:**
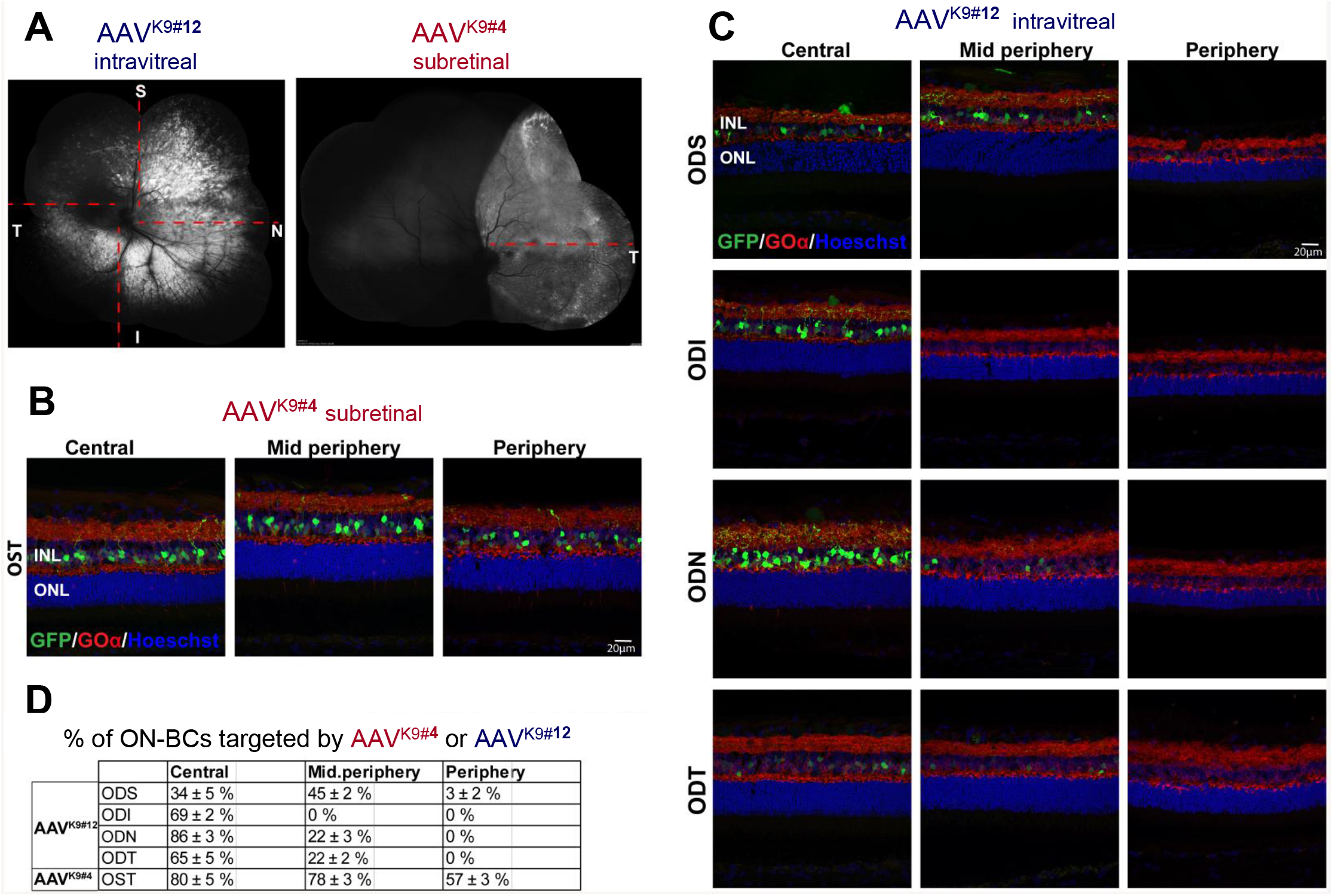
Targeting of canine ON-BCs with top performing AAV capsid variants. **A**) cSLO images (BAF mode) showing GFP fluorescence 6.6 weeks after intravitreal injection of AAV^K9#12^-*lg*GRM6-*sfGFP* (0.38 mL at 2 x10^13^ vg/mL) in the right eye, and subretinal injection of AAV^K9#4^-*lg*GRM6-*sfGFP* (0.15 mL at 3 x10^13^ vg/mL) in the left eye. Dashed red line shows approximate location of retinal cryosections used for IHC and shown in B-C. **B-C**). Retinal cryosections showing native sfGFP fluorescence and Goα immunolabeling. **D**) Percentage of ON-BCs (Goα-positive) that co-express sfGFP transgene in various retinal locations (n=3 technical replicates/eye). OD, right eye; OS, left eye; S, superior; I, inferior; T, temporal; N, nasal; ONL, outer nuclear layer; INL, inner nuclear layer.

To assess applicability of the selected vectors in both subretinal and intravitreal routes of delivery, reporter constructs AAV^K9#4^-*lg*GRM6-*sfGFP* and AAV^K9#12^-*lg*GRM6-*tdTomato* were injected in a cynomolgus macaque. In one eye, both vectors were injected via routes per the canine study whereas injection routes were switched in the contralateral eye (i.e., AAV^K9#4^ intravitreally and AAV^K9#12^ subretinally) (**Fig. S3**). cSLO autofluorescence imaging performed 6 weeks post-injection showed distinct spatial pattern of sfGFP fluorescence between the two eyes. With subretinal injection, both vectors led to transgene expression in ON-BCs throughout the bleb/treated area with a higher transduction rate with AAV^K9#4^ (93%) than with AAV^K9#12^ (80%). Intravitreal injection of either vector led to expression of reporter proteins in ON-BCs in limited areas including the fovea and punctate areas as detected by cSLO. At the level of the fovea/perifovea, transgene expression in ON-BCs was more potent with AAV^K9#12^ than with AAV^K9#4^ when injected intravitreally.

### Development of AAV-*LRIT3* vectors and assessment of safety

To test effective targeting of ON-BCs in the canine CSNB model, therapeutic vectors harboring WT canine *LRIT3* were constructed. For efficient transgene expression in ON-BCs, two ON-BC-specific promoters were selected – the *lg*GRM6 promoter (2.2.kb) used in prior testing with reporter genes, and its abbreviated version *sh*GRM6 (0.7kb), both developed and validated by Lu et al. for increased and specific transduction of ON-BCs compared to the commonly used 200En+SV40p promoter in mouse and NHP retinas^21^. Of the two, the *lg*GRM6 promoter was reported to have greater transduction efficiency but reduced packaging capacity due to its length. To alleviate this challenge, two versions of the expression cassette were constructed. Cassette A utilized the *lg*GRM6 promoter upstream of WT canine *LRIT3* cDNA and polyA signal (**Fig. 2A**). Cassette B instead utilized the *sh*GRM6 promoter, and to stabilize transgene expression, a Woodchuck hepatitis virus posttranscriptional regulatory element (WPRE) was added in between the canine *LRIT3* and polyA signal (**Fig. 2B**). Each expression cassette was then packaged into AAV^K9#4^ and AAV^K9#12^ capsids, generating four AAV vector variants.

**Figure 2.**
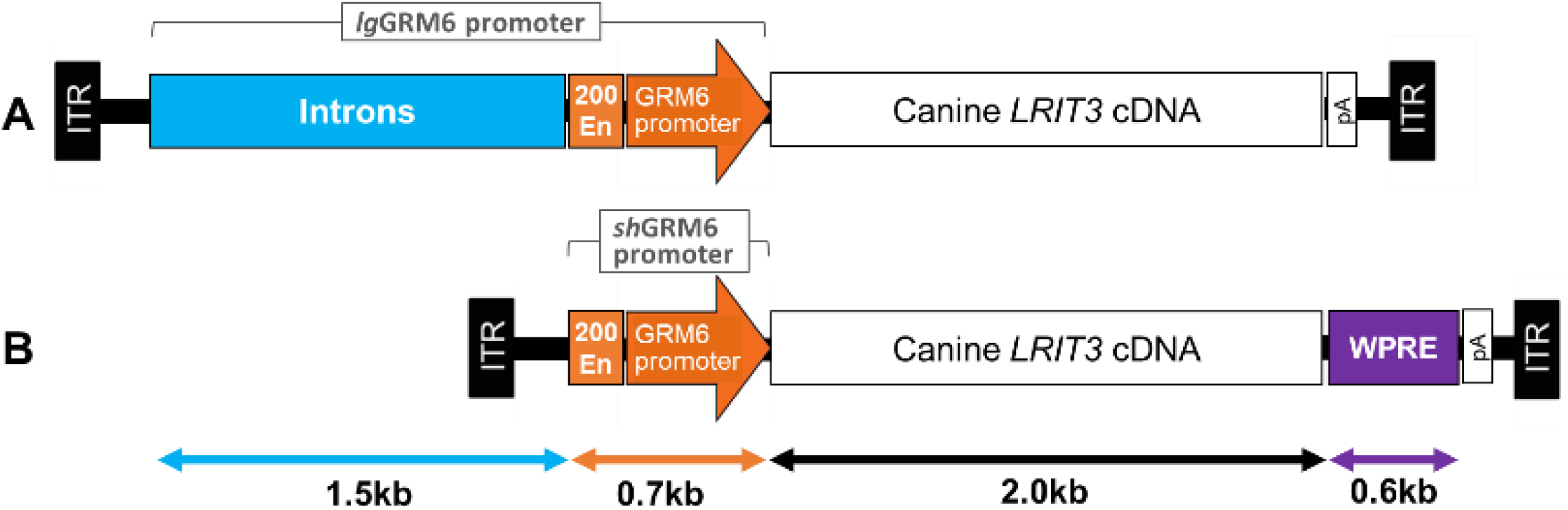
AAV expression cassette variants including GRM6 promoter variants and canine LRIT3 transgene. Schematic diagram of **A**) AAV-*lg*GRM6-c*LRIT3* and **B**) AAV-*sh*GRM6-c*LRIT3*-WPRE cassettes consisting of enhancers (Introns, 200En) and the GRM6 promoter driving canine *LRIT3*. Due to limitations in packaging capacity, vector A lacks WPRE while vector B lacks the Introns and a large part of the enhancers.

In a pilot study for evaluating initial tolerability, each vector was injected in control dogs via the intended route, i.e., AAV^K9#4^ subretinally and AAV^K9#12^ intravitreally, at doses considered high-end for intraocular injections (up to 1×10^13^ vg/mL) (**Table S2**). The two AAV^K9#12^ vectors were administered intravitreally as intended. The two AAV^K9#4^ vectors were administered via the intended subretinal route, although some intravitreal vector leakage occurred during subretinal injections and were recorded. Importantly, no acute or delayed adverse effects were observed for up to 10 weeks post-injection in any of the control eyes as determined by a complete ophthalmic examination including indirect ophthalmoscopy and RetCam® fundus imaging.

### Restoration of ERG b-wave following subretinal AAV^K9#4^-*sh*GRM6-c*LRIT3*-WPRE

To test therapeutic efficacy, the four AAV-*LRIT3* vectors were each injected via the intended route in CSNB dogs (**Table 1**). Of the 8 eyes/4 dogs injected in Cohort 1, two eyes that received subretinal injection of AAV^K9#4^-*sh*GRM6-c*LRIT3*-WPRE (**Fig. 3A**) showed significant scotopic b-wave recovery as early as 8 weeks post-injection, which remained stable at 51 weeks post-injection (**Fig. 3B, Fig. S5A**). Of these two eyes, a more dramatic ERG recovery was observed in the eye that received both subretinal delivery (150 μL) and inadvertent intravitreal leakage (200 μL) (left eye, CHACUG) compared to the eye with subretinal delivery only (left eye, K10) (**Fig. 3B**). None of the other vectors resulted in measurable ERG recovery and were hence deemed “non-therapeutic” and subsequent studies were focused on the “therapeutic” vector (AAV^K9#4^-*sh*GRM6-c*LRIT3*-WPRE).

**Table 1.**
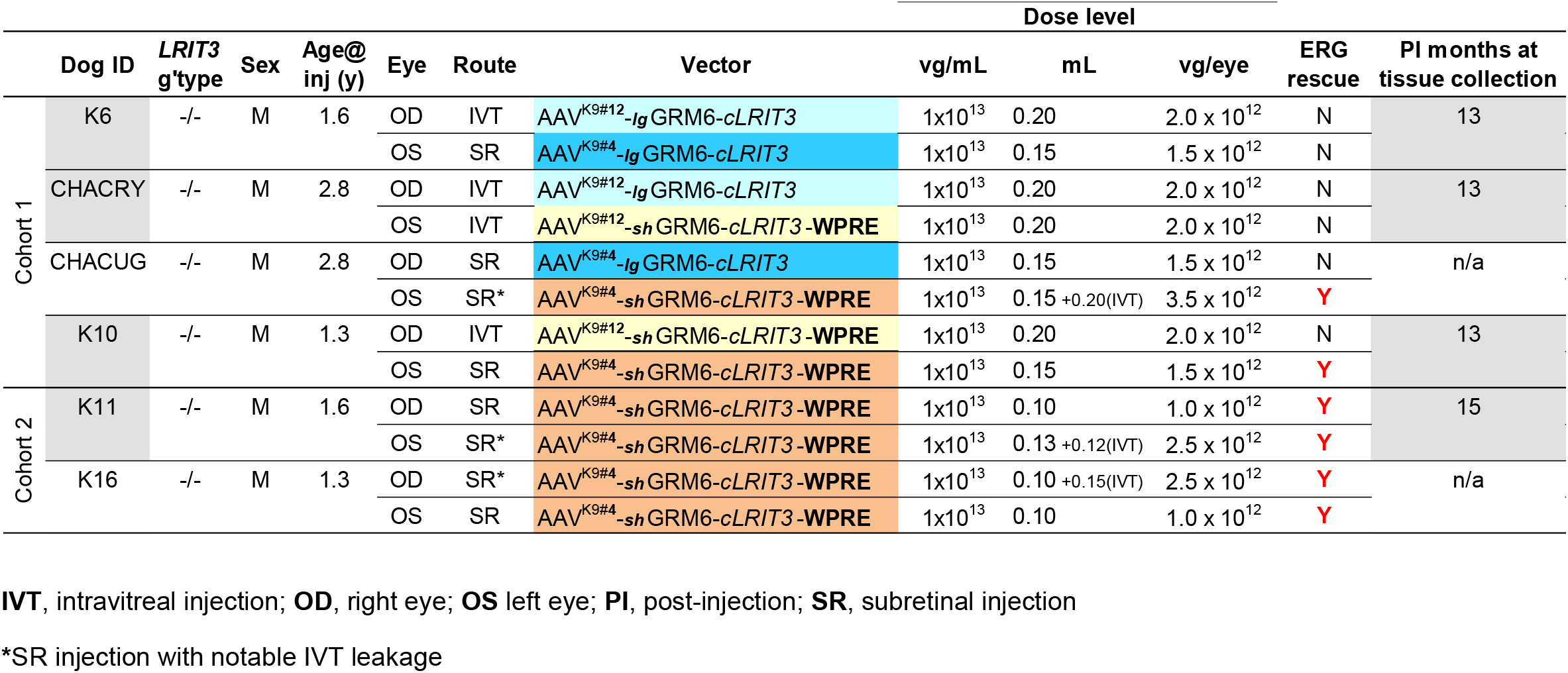
Injection details of efficacy testing in *LRIT3*-CSNB dogs

**Figure 3.**
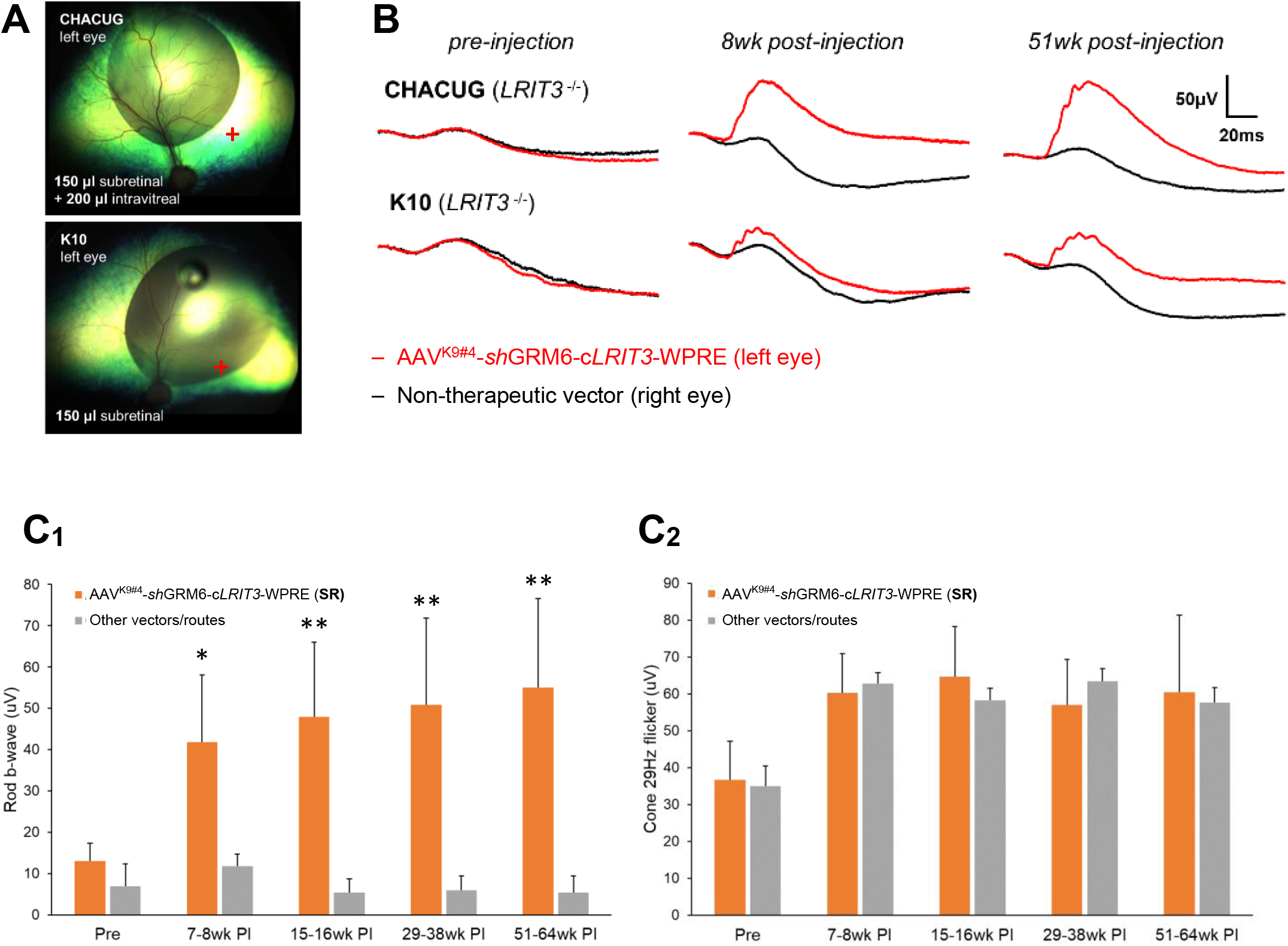
Subretinal AAV-*LRIT3* injection and ERG recovery in CSNB canine eyes. **A**) Subretinal blebs in the left eye of dogs CHACUG and K10 that received subretinal injection of the therapeutic vector AAV^K9#4^-*sh*GRM6-c*LRIT3*-WPRE. Note the injection variables between the two eyes such as bleb location as well as intravitreal vector leakage (200 μL) that occurred in CHACUG. +: canine fovea-like region. **B**) Rod-derived ERG b-wave recovery in treated eyes shown in A). Pre-injection ERG traces demonstrate typical CSNB phenotype with diminished b-wave in both eyes. The eyes receiving subretinal injection of the therapeutic vector show significant b-wave recovery as early as 7 weeks post-injection, with stability up to 64 weeks post-injection. **C**) Rod-derived ERG b-wave (C_1_) and cone-derived 29Hz flicker response (C_2_) pre- and post-injection over time, up to 64 weeks. The eyes subretinally injected with the therapeutic vector (n=6, orange bar) had significant (*p<0.005, **p<0.001) and stable rod-mediated ERG b-wave recovery compared to all other vector/routes (n=6, grey bar) at each post-injection time points. SR, subretinal injection.

Based on the greater b-wave recovery observed with concomitant intravitreal leakage, the effect of combined subretinal and intravitreal injections of the therapeutic vector was tested in Cohort 2. Two dogs received a combined subretinal and intravitreal injection in one eye and subretinal only in the other (**Table 1**). This time, there was no significant difference between the ERG b-wave recovery between the different routes of injection (subretinal + intravitreal vs. subretinal only), both resulting in comparable rod ERG b-wave rescue in one of the two dogs (**Fig. S5B**, K11). Across both cohorts, the eyes subretinally injected with the therapeutic vector had a rod-derived ERG b-wave amplitude that was significantly increased compared to eyes treated with other vectors/routes at each time point. The effect was evident as soon as 7 weeks post-injection (p<0.005) and thereafter remained robust until at least 64 weeks post-injection (p<0.001) (**Fig. 3C**_**1**_). There was no consistent effect on cone-derived ERG associated with specific vectors and injection routes (**Fig. 3C**_**2**_, **Fig. S6**).

### Improved night vision in the eyes treated with subretinal AAV^K9#4^-*sh*GRM6-c*LRIT3*-WPRE

To evaluate the effect of gene therapy on scotopic visual function, injected animals were tested for their ability to navigate in an obstacle avoidance course under scotopic and photopic conditions, for each eye using an occluder. Because CSNB does not affect day vision, testing was done under the three dimmest scotopic conditions (0.003 lux, 0.009 lux, and 0.03 lux) to assess night vision while a photopic condition (65 lux) was used as the control. The CSNB control eye (**Fig. 4, Fig. S7**: white bar) had increased transit times and more collisions at the dimmest light intensity (0.003 lux) as expected from the disease^19^. The phenotype was less prevalent as the light intensity was increased under scotopic conditions (0.009 and 0.03 lux) and normalized at the photopic condition (65 lux). All six CSNB eyes subretinally injected with the therapeutic vector (AAV^K9#4^-*sh*GRM6-c*LRIT3*-WPRE) recovered night vision with significantly faster transit times and fewer collisions at the dimmest light intensity (**Fig. 4, Fig. S7**: solid orange bar) (**Movie S1, S2**) consistent with the observed ERG recovery. Interestingly, eyes subretinally injected with the vector containing the longer promoter (AAV^K9#4^-*lg*GRM6-c*LRIT3*) which did not result in ERG recovery, showed a modest trend of visual rescue at 0.003 lux (**Fig. S7**, solid blue bar). All other vectors injected intravitreally (**Fig. S7**, hatched orange bar) showed no improvement in night vision (**Movie S1, S2**). Strikingly, the recovered night vision was sustained until at least 43 weeks post-injection (**Movie S2**), indicating stability of the therapeutic effect. All CSNB eyes, regardless of the injections, demonstrated normal day vision at 65 lux (**Movie S1, S2**), as expected.

**Figure 4.**
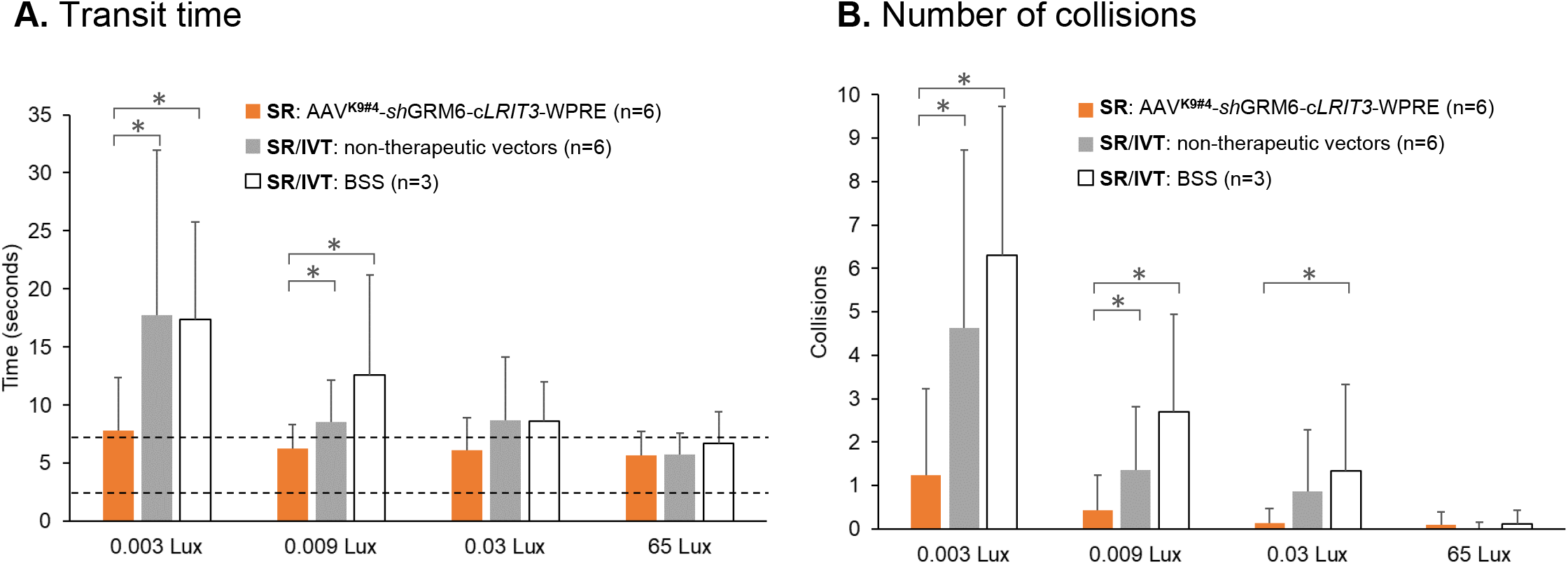
Improved night vision in CSNB canine eyes treated with subretinal AAV^K9#4^-*sh*GRM6-c*LRIT3*-WPRE. Visually-guided navigation tested per eye in an obstacle course in six *LRIT3*-CSNB dogs (12 eyes) injected with one of the AAV vector variants in each eye. **A**) The transit time was significantly reduced following subretinal injection of the therapeutic vector (AAV^K9#4^-*sh*GRM6-c*LRIT3*-WPRE) (6 eyes) compared to the eyes injected with all other vector/routes (6 eyes) or BSS-injected control CSNB eyes at the two dimmest scotopic conditions, 0.003 and 0.009 Lux (*p<.0001). The transition times at 65 Lux (photopic) were minimally affected by the CSNB phenotype and, as such, comparable across different therapeutic and control groups. The dotted lines define the 95% confidence interval for transit time in 4 WT control dogs. **B**) The number of collisions was significantly reduced at 0.003 and 0.009 Lux in eyes subretinally injected with the therapeutic vector (6 eyes) compared to CSNB eyes injected with all other vector/routes (6 eyes) or BSS (*p<.0001). No collisions are observed in WT dogs. While the bars represent the combined data of three available time points 18-50 weeks after the injection, there were no significant changes in vision testing outcomes over time. SR, subretinal; IVT, intravitreal.

### Restoration of LRIT3 in the outer plexiform layer of canine CSNB retina

To localize LRIT3 transgene expression in canine retinas injected with AAV vectors, IHC was performed using antibodies against LRIT3 as well as a variety of pre- and post-synaptic markers. Dogs with the best therapeutic response remain in life and are being followed to assess potential lifetime stability of functional rescue. However, one dog with only modest scotopic b-wave recovery in the eye that was subretinally injected with the therapeutic vector (**Fig. 3A, 3B**; K10, left eye) was sacrificed for immunohistochemical analysis of the retina along with controls. The LRIT3 signals were detected in the outer plexiform layer (OPL) in WT canine retina (**Fig. 5A, D, G**). In this group, LRIT3 labeling appeared to be associated with the synapses between the photoreceptor (PSD95) and second order neurons (Goα) (**Fig 5A**), present in synapses between rods (CtBP2) and rod ON-BCs (PKCα) (**Fig 5D**), and in synapses between cones (HCA) (**Fig 5G**) and all ON-BCs (Goα) (**Fig 5A**). This signal was greatly reduced with the CSNB phenotype^20^. In the eye treated with subretinal injection of the therapeutic vector, within the bleb area, there was limited but distinct LRIT3 signal restored in the OPL, adjacent to both rod (**Fig. 5E**) and cone terminals (**Fig. 5H**). No signal was present outside of the bleb area in the same eye. The canine CSNB eye intravitreally injected with one of the non-therapeutic vectors (AAV^K9#12^-*sh*GRM6-c*LRIT3*-WPRE) (**Fig. S4, S5, S6**; dog CHACRY, left eye) had no evidence of punctate LRIT3 labeling (**Fig. 5C, F, I**). These findings indicate subretinal delivery of the therapeutic vector and expression of the transgene restores LRIT3 to the OPL and contributes to the structural functionality of the synapse by positioning itself to the normal location, at lower levels compared to WT.

**Fig 5.**
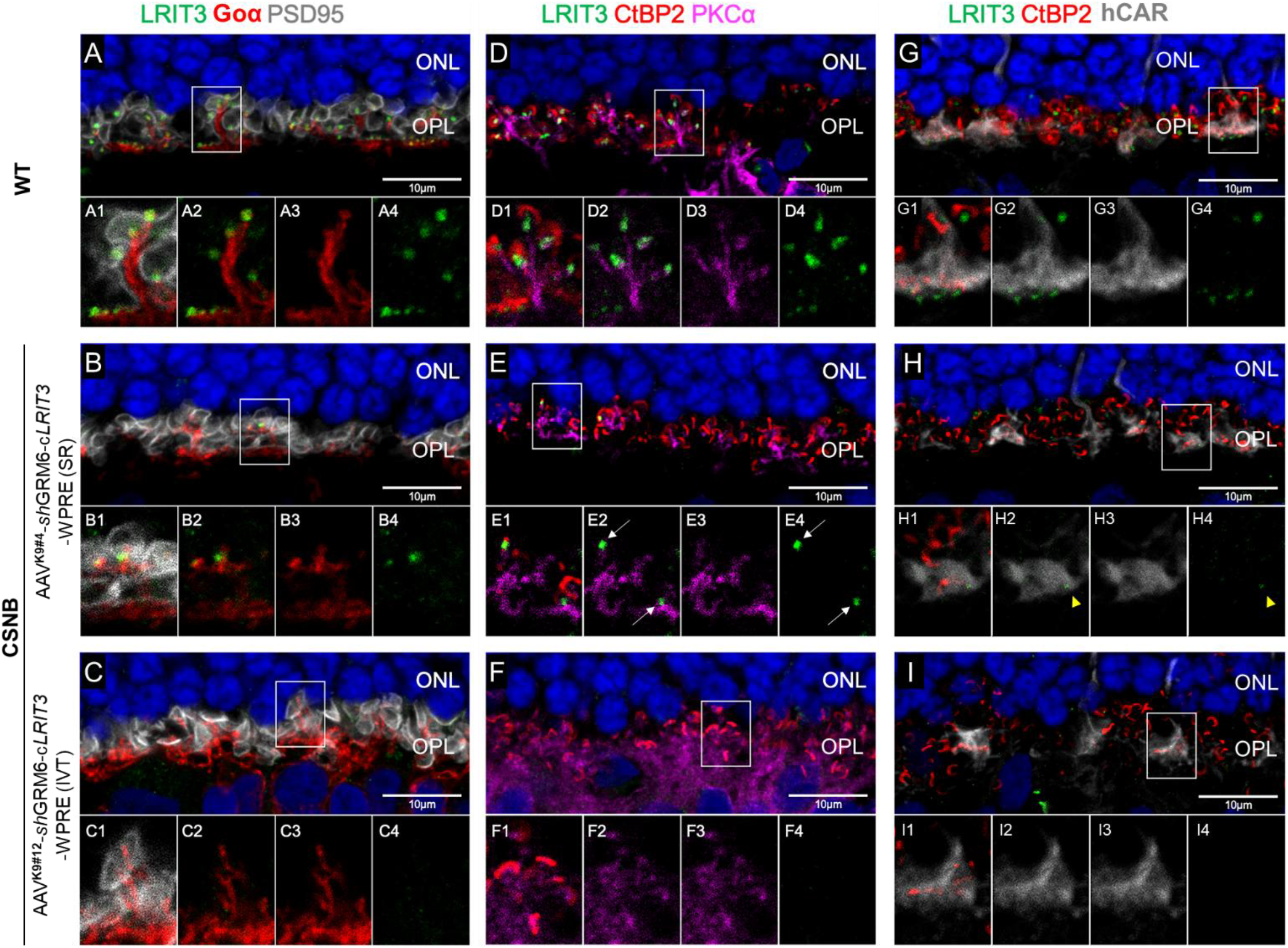
Restoration of LRIT3 in OPL of canine CSNB retina treated with subretinal AAV-*LRIT3*. WT and CSNB canine retinas immunolabelled with LRIT3 together with a variety of pre- and post-synaptic markers. WT canine retina (A, D, G) exhibits punctate LRIT3 signals in OPL adjacent to rod (D1) and cone (G1) terminals. LRIT3 signal is restored, albeit to a limited amount, in the bleb area of canine CSNB retina treated with subretinal AAV-*LRIT3* and showing ERG recovery (B, E, H). Restored LRIT3 signal is present adjacent to the synaptic terminals of both rods (E, arrows) and cones (H, arrowhead). Canine CSNB retina treated with intravitreal AAV-*LRIT3* without ERG recovery (C, F, I) lacked the LRIT3 signals in OPL. IVT, intravitreal; SR, subretinal.

## Discussion

In a newly developed canine model of *LRIT3*-CSNB, we demonstrate safety and efficacy of an AAV gene therapy approach using novel AAV capsid variants and improved mGluR6 promoters, each designed to specifically target ON-BCs. Phenotypic reversal was assessed by ERG and vision-guided behavior revealing functional restoration with 100% efficacy in all six eyes injected subretinally with the therapeutic vector AAV^K9#4^-*sh*GRM6-c*LRIT3*-WPRE, providing excellent proof-of-concept.

We based our approach on previously successful AAV gene therapies delivered to outer retinal cells (i.e., RPE, rod, cone) in canine IRD models^22-28^. Two novel AAV capsid variants that originated from a prior directed evolution screen conducted in WT dogs^29^ were found to specifically target ON-BCs via intravitreal (AAV^K9#12^) or subretinal (AAV^K9#4^) injections in WT dogs. Subsequently, these same variants were shown to effectively transduce ON-BCs in NHP. These new variants constitute novel tools for gene therapy to treat patients with various forms of cCSNB, and for delivery of optogenetic channels to these mid-retinal cell populations in patients with advanced photoreceptor degeneration.

Combining each capsid with two different mGluR6 promoters, we tested four unique vector constructs, and identified a single “therapeutic vector” (AAV^K9#4^-*sh*GRM6-c*LRIT3*-WPRE) candidate. This therapeutic vector, when injected subretinally, consistently led to the significant and stable functional rescue in all six treated eyes. However, there were some noticeable variations in the degree of phenotypic rescue as quantified by recovery of scotopic b-wave amplitude. Possible injection-related factors affecting phenotypic rescue between eyes include inadvertent intravitreal leakage of vector during the subretinal injection procedure and/or location of the subretinal bleb relative to the area centralis.

The hypothesis that a combined subretinal and intravitreal delivery of the therapeutic vector provides superior efficacy to subretinal delivery alone remains to be confirmed. While the therapeutic vector was originally optimized for subretinal injection, its ability to transduce ON-BC via solely the intravitreal route remains to be tested. Nevertheless, in our experience, some degree of intravitreal deposition of vectors while performing subretinal injections is expected, and varying degrees of vector exposure interaction with vitreous humor is inevitable. It is therefore critical that vectors intended for subretinal delivery to be assessed both for toxicity risks and potential transduction benefits resulting from intravitreal leakage.

The current study was conceived prior to the report by Hasan et al. which proposed an alternative, pre-synaptic localization of LRIT3 in the mouse retina^30^. While this localization may be mouse-specific, the gene therapy presented herein was designed for specific targeting of ON-BCs to deliver functional *LRIT3* in the canine *LRIT3*-CSNB model. The AAV vectors used were identified following a screen in WT dogs and selected for ON-BC tropism when administered by subretinal or intravitreal routes. Furthermore, the *sh*GRM6 promoter, derived as a modified version of the mGluR6 (*GRM6*) promoter, was developed and verified by Lu et al. for specific and increased ON-BC expression in mouse and NHP retinas^21^. As well, in our study, subretinal injection of the vector-promoter-reporter constructs in NHP clearly demonstrated ON-BC expression.

While the exact cell of origin and species-specificity of LRIT3 expression in the retina is being resolved, functional rescue from specific targeting of ON-BCs via AAV gene therapy in the canine *LRIT3*-CSNB is striking and sheds light into the cellular origin as well as the optimal functional location of LRIT3. Hasan et al. reported ERG recovery in *Lrit3*^-/-^ mice receiving AAV-*Lrit3* targeted to the rod photoreceptors, but this was successful only with early treatment at P5, and with limited effect with late treatment at P30 (**Table S1**). Interestingly there was rod-specific transgene expression following the injections at either timepoints^30^, suggesting that physical augmentation of LRIT3 in rods may not always be capable of restoring function in mature retina, thus limiting its translational potential. In a recent report by Varin et al., adult *Lrit3*^nob6^ mice were treated intravitreally using AAV2-7m8-*Lrit3*, targeting either ON-BCs or photoreceptors alone, or both (OPL) by interchanging the cell-specific promoters^31^. Restoration of LRIT3 and TRPM1 localization, and functional rescue were demonstrated in only a fraction (7-25%) of the treated mice (**Table S1**). While targeting of either or both cells resulted in a few responsive animals, rate and degree of functional recovery was greater with targeting the photoreceptors, followed by OPL and then ON-BCs, again pointing to potential species-specific differences. In our current work, functional recovery was achieved in adult *LRIT3*-CSNB dogs treated at ages ranging from 1.3 to 2.8 years, demonstrating therapeutic efficacy in mature retina with the CSNB phenotype in 100% (6/6) of eyes injected subretinally with the therapeutic vector targeting ON-BCs. Future work will evaluate if efficacy may be improved if *LRIT3*-CSNB animals are treated at a younger age, and if augmentation therapy is effective in animals older than 3 years of age.

The therapeutic vector was the only one of four vectors tested resulting in scotopic ERG b-wave functional rescue. However, of the three non-therapeutic vectors, AAV^K9#4^-*lg*GRM6-c*LRIT3* injected subretinally in two dogs demonstrated modest improvement in scotopic vision-guided behavior parameters compared to controls despite having no detectable recovery of rod b-wave function. This indicates the *lg*GRM6-c*LRIT3* expression cassette may be functional albeit less efficient than the *sh*GRM6-c*LRIT3*-WRPE cassette. In contrast, Lu et al. reported that the *lg*GRM6 promoter was more efficient than *sh*GRM6^21^, suggesting that other factors could affect the expression efficiency in our study, such as improved expression driven by WPRE or interference due to the expression cassette nearing maximum capacity, which was the case for *lg*GRM6-c*LRIT3*.

The mildly improved scotopic function in the eye receiving subretinal AAV^K9#4^-*lg*GRM6-c*LRIT3* was only detectable by a vision-guided obstacle course navigation and not by ERG, indicating that the former outcome measure is more sensitive for detecting minor functional recovery with potential clinical relevance. This is consistent with the previous assessment of therapeutic outcome in Leber congenital amaurosis (LCA) patients receiving AAV therapy^32,33^, and re-emphasizes the importance of incorporating intuitive and straightforward vision-guided behavior as an outcome assessment of therapy. None of the vectors that utilized the AAV^K9#12^ capsid designed for intravitreal injection were therapeutically effective. While the initial screening of this AAV capsid variant was done in intact canine eyes, going forward pre-treatment of diseased eyes with vitrectomy or enzymatic vitreolysis^34^ may improve access of the vector to the retinal layers.

We hereby demonstrate safety and efficacy of a novel AAV gene therapy designed to target ON-BCs in the canine *LRIT3*-CSNB model. Significant and consistent functional recovery was achieved up to 1.2 years post-injection as assessed by ERG and vision-guided behavior testing. This is the first patient-relevant data showing rescue of ON-BC function by gene augmentation therapy using a translationally suitable large animal model of *LRIT3*-CSNB. With intact retinal architecture and reliable and validated outcome measures available, the canine *LRIT3*-CSNB model has translational applicability for determination of optimal treatment strategy for CSNB patients. In addition, the model will serve more broadly as a platform to test therapeutic products which target OPL. As the evolving narrative for cell of origin of native LRIT3 in mature retina is further refined, it will guide our quest to identify and test the most optimal therapeutic targets for the treatment of *LRIT3*-CSNB.

## Materials & Methods

### Ethics statement

The research was conducted in strict accordance with the recommendations in the Guide for the Care and Use of Laboratory Animals of the National Institutes of Health, in full compliance with the USDA’s Animal Welfare Act, Animal Welfare Regulations, and the ARVO Statement for the Use of Animals in Ophthalmic and Vision Research. The protocols were approved by the Institutional Animal Care and Use Committee of the University of Pennsylvania (IACUC# 803269), and Charles River Laboratories.

### Animals

The *LRIT3*-CSNB model originated from a research colony that was used for disease characterization^19^ and mutation identification^20^. The dogs in the current study are part of a newly established *LRIT3*-CSNB research line developed at the Retinal Disease Studies Facility (RDSF) of the University of Pennsylvania, and formed by outcrossing dogs from the original colony^19^ to WT dogs and to unrelated dogs that had been identified to harbor the *LRIT3* variant through molecular screening. Both control (WT and *LRIT3*-CSNB carrier) and affected dogs, including an *NPHP5*-LCA dog used in AAV vector screening, were bred at the RDSF or by a commercial laboratory animal supplier and maintained and housed at the RDSF under identical conditions of diet, medications, vaccinations, and ambient illumination with cyclic 12h ON-12h OFF. CSNB affected dogs were identified based on genotyping of the *LRIT3* genetic variant^20^ and the characteristic loss of scotopic ERG b-wave^19^. A single adult male cynomolgus macaque housed at Charles River Laboratories, Mattawan, MI, was used in this study to validate the ON-BC tropism of top performing variants identified following a screen in WT dogs.

### Promoter and AAV capsid screen for bipolar cell targeting

Eight different AAV capsid variants originating from an AAV2-7-mer library that had undergone directed-evolution screens in dogs^29^ for their ability to target the outer retina after intravitreal injection were selected together with AAV2_4YF+TV_ (used as a control) for this current study (**Table S3**). Each of these 9 capsids were used individually to package a recombinant genome containing the ON-BC-specific promoter Ins4s-In3-200En-mGluR500p (named thereafter *lg*GRM6)^21^ driving expression of sfGFP followed by a 3’-barcode unique to that capsid. The vectors were diluted until the titers of all variants were equal, as confirmed by quantitative real-time PCR, and then combined in equal ratios by adding equivalent volume of each vector solution to a pool. A similar approach was used to produce a second pool containing the same nine AAV capsids carrying sfGFP cDNA with the unique barcode but under control of the 4xGRM6^35^ ON-BC-specific promoter. A third pool of the same AAVs was also made but with the Ple155^36^ ON-BC-specific promoter. The latter two pools showed lack of ON-BC specificity when injected intravitreal or subretinal with AAV2_4YF+TV_ vector and were not developed further for this study.

### GFP-barcoded library screening in canine retina

Seven weeks following subretinal (1 eye) and intravitreal (2 eyes) injections in two WT dogs, biopsies of neuroretina taken from eyes enucleated after humane euthanasia were collected from locations in which sfGFP fluorescence was detected by cSLO. RNA was purified from these samples and deep sequencing was performed to quantify the relative extents to which each capsid was capable of delivering and expressing its transgene in the canine neuroretina from both the vitreous and subretinal space. Quantification of AAV capsid performance revealed that AAV^K9#12^ and AAV^K9#4^ outperformed other variants when delivered via the intravitreal and subretinal routes, respectively, and thus were selected for further validation.

### AAV-*LRIT3* viral constructs

To identify the most optimal expression cassette for robust and stable therapeutic gene expression, AAV expression cassette variants *lg*GRM6-c*LRIT3* and *sh*GRM6-c*LRIT3*-WRPE were constructed (**Fig. 2**). In both variants, the therapeutic gene consisted of full-length canine *LRIT3* (c*LRIT3*) cDNA, with a canonical Kozak sequence incorporated at the translational initiation site. The c*LRIT3* sequence was cloned from WT canine retinal cDNA, verified by Sanger sequencing, and corrected for polymorphic changes in amino acids to match those of the canine reference genome (CamFam3.1) and of the consensus across mammalian species. Improved mGluR6 promoters developed by Lu et al. aimed at targeting ON-BCs^21^ were provided by Dr. Z-H Pan of Wayne State University. The c*LRIT3* transgene was placed under the control of either the original full-length promoter (In4s-In3-200En-mGluR500P^21^, *lg*GRM6) or the shortened promoter (200En-mGluR500P^21^, *sh*GRM6). For transcript stabilization, a WPRE sequence was added downstream of the transgene with the *sh*GRM6 promoter. This was not possible with the *lg*GRM6 promoter due to packaging capacity limitations. The two construct variants were each packaged into AAV^K9#4^ and AAV^K9#12^ capsids selected via directed-evolution for subretinal and intravitreal injections, respectively, as described above. AAV vectors were produced by triple plasmid co-transfection purified and titered as previously described^37^.

### Vector administration and post-injection treatment

Under general anesthesia, viral vectors were delivered by subretinal or intravitreal injections in control and affected canine eyes (**Table S2** and **Table 1** show animals, vectors, and injection details in control and CSNB dogs, respectively), as well as in a single NHP. Viral vectors were diluted in sterile balanced salt solution (BSS) and delivered 100-200 μL per eye. A transvitreal approach was used without vitrectomy via *pars plana* by using a custom-modified subretinal injector with a 39-gauge polyimide cannula (Retinaject; Surmodics, Inc., Eden Prairie, MN)^38,39^ under direct visualization through Machemer magnifying lens (OMVI; Ocular Instruments Inc., Bellevue, WA, USA) using an operating microscope. Subretinal delivery was directed at the superotemporal quadrant aiming to form a large uniform bleb, encompassing approximately 20-40% of the retina (**Fig S4**), which was immediately visualized and documented by fundus photography (Retcam shuttle; Natus Medical, Inc., Pleasanton, CA). Intravitreal delivery was performed by deposition of viral solution in the posterior vitreous just pre-retinally at four sites horizontally across the visual streak. Anterior chamber paracentesis was performed immediately post-injection to prevent increase in intraocular pressure. The post-injection antibiotic and anti-inflammatory routine treatment plan has been described previously^23^.

### Ophthalmic examination

Ophthalmic examinations included slit-lamp biomicroscopy, tonometry, indirect ophthalmoscopy, and fundus photography. Any abnormalities, including any development of inflammation or lesions, were documented. The ophthalmic examinations were conducted pre-injection, 1- and 2-day post-injection, weekly for the following 4 weeks, bi-weekly for additional 8 weeks, then every 12-16 weeks up to 64 weeks post-injection.

### Electroretinography

Recordings were conducted at pre-injection, then post-injection time points of 7-8, 15-16, 29-38, and 51-64 weeks after vector delivery. Pupils were dilated with topical atropine sulfate 1% (Akorn, Inc., Lake Forest, IL), tropicamide 1% (Akorn, Inc.), and phenylephrine 10% (Paragon Biotech, Portland, OR). After induction with intravenous propofol (Zoetis, Kalamazoo, MI), dogs were maintained under general inhalation anesthesia (isoflurane 2-3%; Akorn, Inc.). Full-field flash electroretinography (ERG) was performed on both eyes using a custom-built Ganzfeld dome fitted with LED stimuli from a ColorDome stimulator (Diagnosys LLC, Lowel, MA). After 20 min of dark adaptation, rod and mixed rod-cone-mediated responses to single 4 ms white flash stimuli of increasing intensities (−3.7 to 0.5 log cd.s.m^-2^) were recorded. After 5 min of white light adaptation (10.6 cd.m^-2^), cone-mediated responses to a series of single white flashes (−2.7 to 0.5 log cd.s.m^-2^) and to 29.4 Hz flicker stimuli (−2.7 to 0.2 log cd.s.m^-2^) were recorded. Finally, photopic long-flash ERGs were recorded using 200 msec white stimuli of 400 cd.m^-2^ on a rod-suppressing white background of 10.6 cd.m^-2^ to assess the ON- and OFF-pathways. Statistical significance was calculated by an unpaired t-test using GraphPad (GraphPad Software, San Diego, CA).

### Obstacle course vision testing

Vision-guided navigation was tested in an obstacle avoidance course as previously described^40^. An abbreviated protocol was used for this study under the three dimmest scotopic conditions (0.003 lux, 0.009 lux, and 0.03 lux) and under ambient photopic condition (65 lux, fluorescent room light). Each eye was tested individually by having an Aestek opaque corneoscleral shield (Oculo-plastik Inc., Montreal, Canada) placed over the ocular surface of the contralateral eye after topical anesthesia (proparacaine 0.5%). All eyes were tested three times under each light intensity, and the position of the 5 panels was randomly changed between each of the 3 trials/eye/illumination. The contralateral eye was tested with the same set of panel positions. The eye to be tested was randomly selected before the session. Animals were first dark-adapted for 20 min to the lowest ambient illumination (0.003 lux) before running through the course. After testing under scotopic conditions, room illumination was increased to the photopic brightness to light-adapt animals for 10 min and animals were tested under photopic conditions in the same manner. Two digital Sony Handycam DCR-DVD108 cameras (Sony, San Diego, CA) located above the obstacle course recorded navigation performance of the dogs. The infrared imaging function of the camera enabled recording under the dimmest light conditions. An experienced observer who was masked to experimental design reviewed all videos and measured for each trial total number of collisions and transit time in seconds between first forward motion at course entrance and moment the animal completely passed through the exit gate. Vision testing was repeated at least three times on different dates post-injection to ensure inter-testing consistency. An unpaired t-test was carried out using GraphPad (GraphPad Software, San Diego, CA) to analyze statistical significance in number of collisions and transit time under each illumination level between groups with a cut-off of p<0.0001.

### Immunohistochemistry (IHC)

Preparation and processing of tissues for retinal morphology and IHC have been detailed previously^20,23^. Antibodies used included GFP (Millipore cat# MAB3580; Abcam cat# ab290), tdTomato (SICGEN cat# AB8181-200), LRIT3 (Sigma-Aldrich, cat# HPA013454), pre-synaptic markers PSD95 (BioLegend cat#810401), goat anti-human CAR (custom) and CtBP2 (BD Biosciences, cat# 612044), and post-synaptic markers PKCα (BD Biosciences, cat# 610107) and Goα (Millipore cat# MAB3073). Slides were examined with a Nikon A1R confocal microscope (Nikon Instruments Inc., Melville, NY). Digital images were captured and processed using the NIS Elements (Nikon Instruments Inc., Melville, NY) and ImageJ software (National Institutes of Health).

## Acknowledgements

We are grateful to Dr. Mineo Kondo of Mie University, Mr. Kazuhiko Sakai and Mr. Takehiro Aihara of Kitayama Labes, Japan, for providing foundation dogs for the establishment of the canine CSNB research colony. Additional foundation dogs were generous donations from Labcorp Drug Development, formerly Covance. The *lg*GRM6 and *sh*GRM6 promoters were kindly provided by Dr. Zhuo-Hua Pan of Wayne State University. We also thank Ms. Courtney Spector, formerly of University of Pennsylvania for technical assistance, Dr. András Komáromy, Michigan State University, for procedural instructions, Dr. Leslie King, University of Pennsylvania, for critical review of the manuscript. This study was supported, in part, by Margaret Q. Landenberger Research Foundation (KM), NEI/NIH grant EY-006855 (GDA/KM), Foundation Fighting Blindness (GDA/WB), the Van Sloun Foundation for Canine Genetic Research (GDA), and the Sanford and Susan Greenberg End Blindness Outstanding Achievement Prize (GDA).

## References

1. Koike C, Obara T, Uriu Y, et al. TRPM1 is a component of the retinal ON bipolar cell transduction channel in the mGluR6 cascade. Proc Natl Acad Sci U S A 2010;107:332–337.

2. Neuille M, Morgans CW, Cao Y, et al. LRIT3 is essential to localize TRPM1 to the dendritic tips of depolarizing bipolar cells and may play a role in cone synapse formation. Eur J Neurosci 2015;42:1966–1975.

3. Zeitz C, Robson AG, Audo I. Congenital stationary night blindness: an analysis and update of genotype-phenotype correlations and pathogenic mechanisms. Prog Retin Eye Res 2015;45:58–110.

4. Riggs LA. Electroretinography in cases of night blindness. Am J Ophthalmol 1954;38:70–78.

5. Schubert G, Bornschein H. [Analysis of the human electroretinogram]. Ophthalmologica 1952;123:396–413.

6. Miyake Y, Yagasaki K, Horiguchi M, et al. Congenital stationary night blindness with negative electroretinogram. A new classification. Arch Ophthalmol 1986;104:1013–1020.

7. Bech-Hansen NT, Naylor MJ, Maybaum TA, et al. Loss-of-function mutations in a calciumchannel alpha1-subunit gene in Xp11.23 cause incomplete X-linked congenital stationary night blindness. Nat Genet 1998;19:264–267.

8. Pusch CM, Zeitz C, Brandau O, et al. The complete form of X-linked congenital stationary night blindness is caused by mutations in a gene encoding a leucine-rich repeat protein. Nat Genet 2000;26:324–327.

9. Audo I, Kohl S, Leroy BP, et al. TRPM1 is mutated in patients with autosomal-recessive complete congenital stationary night blindness. Am J Hum Genet 2009;85:720–729.

10. Li Z, Sergouniotis PI, Michaelides M, et al. Recessive mutations of the gene TRPM1 abrogate ON bipolar cell function and cause complete congenital stationary night blindness in humans. Am J Hum Genet 2009;85:711–719.

11. van Genderen MM, Bijveld MM, Claassen YB, et al. Mutations in TRPM1 are a common cause of complete congenital stationary night blindness. Am J Hum Genet 2009;85:730–736.

12. Dryja TP, McGee TL, Berson EL, et al. Night blindness and abnormal cone electroretinogram ON responses in patients with mutations in the GRM6 gene encoding mGluR6. Proc Natl Acad Sci U S A 2005;102:4884–4889.

13. Zeitz C, van Genderen M, Neidhardt J, et al. Mutations in GRM6 cause autosomal recessive congenital stationary night blindness with a distinctive scotopic 15-Hz flicker electroretinogram. Invest Ophthalmol Vis Sci 2005;46:4328–4335.

14. Audo I, Bujakowska K, Orhan E, et al. Whole-exome sequencing identifies mutations in GPR179 leading to autosomal-recessive complete congenital stationary night blindness. Am J Hum Genet 2012;90:321–330.

15. Peachey NS, Ray TA, Florijn R, et al. GPR179 is required for depolarizing bipolar cell function and is mutated in autosomal-recessive complete congenital stationary night blindness. Am J Hum Genet 2012;90:331–339.

16. Zeitz C, Jacobson SG, Hamel CP, et al. Whole-exome sequencing identifies LRIT3 mutations as a cause of autosomal-recessive complete congenital stationary night blindness. Am J Hum Genet 2013;92:67–75.

17. Neuille M, El Shamieh S, Orhan E, et al. Lrit3 deficient mouse (nob6): a novel model of complete congenital stationary night blindness (cCSNB). PLoS ONE 2014;9:e90342.

18. Neuille M, Cao Y, Caplette R, et al. LRIT3 Differentially Affects Connectivity and Synaptic Transmission of Cones to ON- and OFF-Bipolar Cells. Invest Ophthalmol Vis Sci 2017;58:1768–1778.

19. Kondo M, Das G, Imai R, et al. A Naturally Occurring Canine Model of Autosomal Recessive Congenital Stationary Night Blindness. PLoS One 2015;10:e0137072.

20. Das RG, Becker D, Jagannathan V, et al. Genome-wide association study and whole-genome sequencing identify a deletion in LRIT3 associated with canine congenital stationary night blindness. Sci Rep 2019;9:14166.

21. Lu Q, Ganjawala TH, Ivanova E, et al. AAV-mediated transduction and targeting of retinal bipolar cells with improved mGluR6 promoters in rodents and primates. Gene Ther 2016;23:680–689.

22. Acland GM, Aguirre GD, Ray J, et al. Gene therapy restores vision in a canine model of childhood blindness. Nat Genet 2001;28:92–95.

23. Aguirre GD, Cideciyan AV, Dufour VL, et al. Gene Therapy Reforms Photoreceptor Structure and Restores Vision in NPHP5-associated Leber Congenital Amaurosis. Mol Ther 2021.

24. Beltran WA, Cideciyan AV, Lewin AS, et al. Gene therapy rescues photoreceptor blindness in dogs and paves the way for treating human X-linked retinitis pigmentosa. Proc Natl Acad Sci U S A 2012;109:2132–2137.

25. Cideciyan AV, Sudharsan R, Dufour VL, et al. Mutation-independent rhodopsin gene therapy by knockdown and replacement with a single AAV vector. Proc Natl Acad Sci U S A 2018;115:E8547–E8556.

26. Guziewicz KE, Cideciyan AV, Beltran WA, et al. BEST1 gene therapy corrects a diffuse retina-wide microdetachment modulated by light exposure. Proc Natl Acad Sci U S A 2018;115:E2839–E2848.

27. Komaromy AM, Alexander JJ, Rowlan JS, et al. Gene therapy rescues cone function in congenital achromatopsia. Hum Mol Genet 2010;19:2581–2593.

28. Ye GJ, Komaromy AM, Zeiss C, et al. Safety and Efficacy of AAV5 Vectors Expressing Human or Canine CNGB3 in CNGB3-Mutant Dogs. Hum Gene Ther Clin Dev 2017;28:197–207.

29. Öztürk BE, Johnson ME, Kleyman M, et al. scAAVengr: Single-cell transcriptome-based quantification of engineered AAVs in non-human primate retina. BioRxiv 2020.

30. Hasan N, Pangeni G, Cobb CA, et al. Presynaptic Expression of LRIT3 Transsynaptically Organizes the Postsynaptic Glutamate Signaling Complex Containing TRPM1. Cell Rep 2019;27:3107–3116 e3103.

31. Varin J, Bouzidi N, Gauvain G, et al. Substantial restoration of night vision in adult mice with congenital stationary night blindness. Mol Ther Methods Clin Dev 2021;22:15–25.

32. Chung DC, McCague S, Yu ZF, et al. Novel mobility test to assess functional vision in patients with inherited retinal dystrophies. Clin Exp Ophthalmol 2018;46:247–259.

33. Russell S, Bennett J, Wellman JA, et al. Efficacy and safety of voretigene neparvovec (AAV2-hRPE65v2) in patients with RPE65-mediated inherited retinal dystrophy: a randomised, controlled, open-label, phase 3 trial. Lancet 2017;390:849–860.

34. Stalmans P, Benz MS, Gandorfer A, et al. Enzymatic vitreolysis with ocriplasmin for vitreomacular traction and macular holes. N Engl J Med 2012;367:606–615.

35. Cronin T, Vandenberghe LH, Hantz P, et al. Efficient transduction and optogenetic stimulation of retinal bipolar cells by a synthetic adeno-associated virus capsid and promoter. EMBO Mol Med 2014;6:1175–1190.

36. de Leeuw CN, Korecki AJ, Berry GE, et al. rAAV-compatible MiniPromoters for restricted expression in the brain and eye. Mol Brain 2016;9:52.

37. Flannery JG, Visel M. Adeno-associated viral vectors for gene therapy of inherited retinal degenerations. Methods Mol Biol 2013;935:351–369.

38. Beltran WA, Boye SL, Boye SE, et al. rAAV2/5 gene-targeting to rods:dose-dependent efficiency and complications associated with different promoters. Gene Ther 2010;17:1162–1174.

39. Komaromy AM, Varner SE, de Juan E, et al. Application of a new subretinal injection device in the dog. Cell Transplant 2006;15:511–519.

40. Garcia MM, Ying GS, Cocores CA, et al. Evaluation of a behavioral method for objective vision testing and identification of achromatopsia in dogs. Am J Vet Res 2010;71:97–102.

