## Supplemental Tables Figs for "Targeting ON-bipolar cells by AAV gene therapy stably reverses *LRIT3*-congenital stationary night blindness"

**Suppl. Table S1.** Comparison of CSNB disease and animal models with *LRIT3* deficits and outcome of AAV gene therapy preclinical trials.

| Species | Human | Dog | Mouse | | Mouse | | |
|---|---|---|---|---|---|---|---|
| **Disease/Model** | *LRIT3*-CSNB | *LRIT3*-CSNB | *Lrit3*$^{-/-}$ | | *Lrit3*$^{nob6}$ | | |
| **Rod ERG b-wave** | Diminished | Diminished | Diminished | | Diminished | | |
| **Cone ERG b-wave** | **Normal - mildly reduced** | **Normal - mildly reduced** | Diminished | | Diminished | | |
| **Target cells by design** | - | ON-BC | Rods | | ON-BC | PR | OPL |
| **AAV serotype** | - | AAV$^{K9\#4}$ | rAAV2/2[MAX] | | AAV2.7m8 | | |
| **Promoter** | - | *sh*GRM6* | *RHO* | | *Grm6*† | h*GRK* | *Grm6*† & h*GRK* |
| **Admin. route** | - | subretinal | intravitreal | | Intravitreal | | |
| **Age at therapy** | | **adult** (1.3-2.8 yrs) | premature (P5) | adult (P35) | adult (P30) | | |
| **Scotopic b-wave recovery** | - | 6/6 **(100%)** | moderate | limited | 3/43 (7%) | 6/24 (25%) | 1/10 (10%) |
| **Therapeutic duration** | | 64+ weeks | 8 weeks | | 16 weeks | | |
| **Reference** | Miyake et al. 1986 | current study | Hasan et al. 2019 | | Varin et al. 2021 | | |

*Developed by Lu et al. as 200En-mGluR500P
†200bp enhancer

**Suppl. Table S2**: Injection details of safety testing in control dogs.

| | | | | | | | Dose level | | | | |
|---|---|---|---|---|---|---|---|---|---|---|---|
| **Dog ID** | ***LRIT3* g'type** | **Sex** | **Age@ inj. (y)** | **Eye** | **Route** | **Vector** | **vg/mL** | **mL** | **vg/eye** | **Toxicity** | **Post-inj. follow-up** |
| K9 | +/- | F | 1.3 | OD | SR* | AAV$^{K9\#4}$-***sh***GRM6-*cLRIT3*-**WPRE** | $1x10^{13}$ | 0.10 +0.05(IVT) | $1.5 \times 10^{12}$ | N | 10 wks |
| | | | | OS | SR* | AAV$^{K9\#4}$-***lg***GRM6-*cLRIT3* | $1x10^{13}$ | 0.03 +0.12(IVT) | $1.5 \times 10^{12}$ | N | |
| N305 | +/- | M | 2.8 | OD | IVT | AAV$^{K9\#12}$-***sh***GRM6-*cLRIT3*-**WPRE** | $1x10^{12}$ | 0.20 | $2.0 \times 10^{11}$ | N | 8 wks |
| | | | | OS | IVT | AAV$^{K9\#12}$-***sh***GRM6-*cLRIT3*-**WPRE** | $1x10^{13}$ | 0.20 | $2.0 \times 10^{12}$ | N | |
| CBBCDF | +/- | F | 8.7 | OD | IVT | AAV$^{K9\#12}$-***lg***GRM6-*cLRIT3* | $1x10^{12}$ | 0.20 | $2.0 \times 10^{11}$ | N | 8 wks |
| | | | | OS | IVT | AAV$^{K9\#12}$-***lg***GRM6-*cLRIT3* | $1x10^{11}$ | 0.20 | $2.0 \times 10^{10}$ | N | |

**IVT**, intravitreal injection; **SR**, subretinal injection

*Intended as SR injection although unintentional IVT vector leakage occurred at the dose indicated.

**Suppl. Table S3**: Results of AAV capsid variant screen in WT canine retina following intravitreal and subretinal injection.

| INTRAVITREAL | | | SUBRETINAL | | |
|---|---|---|---|---|---|
| ***Capsid variant*** | ***Name*** | ***Score*** | ***Capsid variant*** | ***Name*** | ***Score*** |
| LAPDSTTRSA | K9#12 | 5.36 | LATTSQNKPA | K9#4 | 5.72 |
| LALGETTRPA | 7mer8 | 4.72 | LAKSDQSKPA | K9#5 | 2.75 |
| LAVDGAQRSA | K9#11 | 2.67 | LAPDSTTRSA | K9#12 | 1.48 |
| LATTSQNKPA | K9#4 | 2.35 | LALGETTRPA | 7mer8 | 1.42 |
| LAKSDQSKPA | K9#5 | 1.40 | PAPQDTTKKA | K9#16 | 1.13 |
| PAPQDTTKKA | K9#16 | 0.48 | LAVDGAQRSA | K9#11 | 0.94 |
| $AAV2_{4YF+TV}$ | control | 0.09 | $AAV2_{4YF+TV}$ | control | 0.61 |
| PAHQDTTKNA | K9#6 | 0.04 | LAKDATKTIA | K9#9 | 0.39 |
| LAKDATKTIA | K9#9 | 0.03 | PAHQDTTKNA | K9#6 | 0.38 |

Note: AAV capsid variants are ordered from best (top) to worst (bottom) performing vectors. Scores indicate % of total of recovered library / % total in total AAV library. Rankings were calculated from mRNA recovered from canine retina.

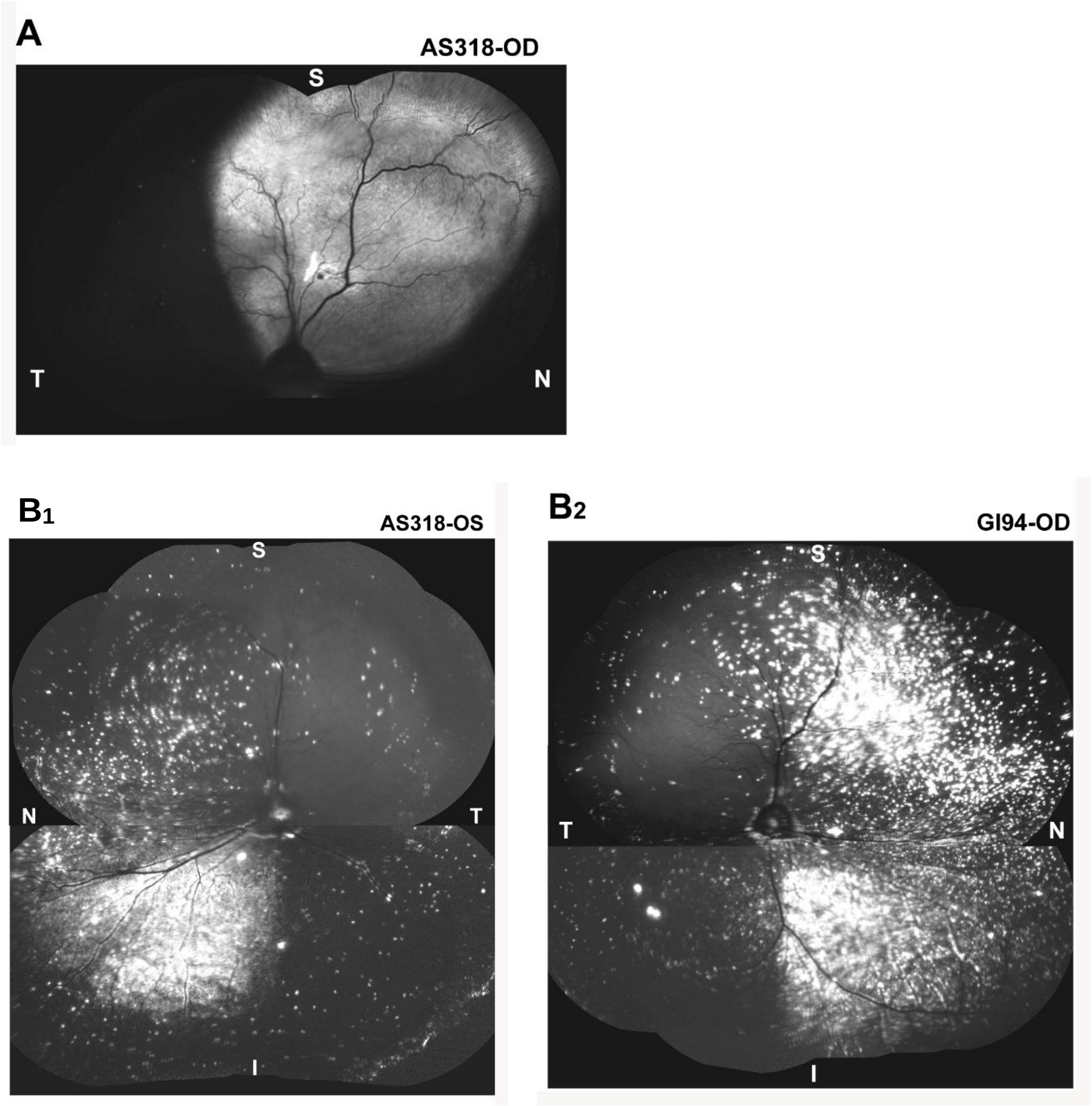


**Suppl. Fig. S1: Detection of GFP fluorescence by cSLO imaging following subretinal and intravitreal injections of a GFP-barcoded mix of AAV capsid variants.**

**A**) Composite cSLO image (BAF mode) 6.4 weeks following subretinal injection (vol: 0.15 ml; titer: 3.9 x $10^{13}$ vg/mL) in the eye of a WT dog (AS318). **$B_1$, $B_2$**) Composite cSLO images 6.4 weeks following intravitreal injection (vol: 0.39-0.4 ml; titer: 3.9 x $10^{13}$ vg/mL) in eyes of two WT dogs (AS318 and GI94). OD, right eye; OS, left eye; T, temporal; N, nasal; T, superior; I, inferior.

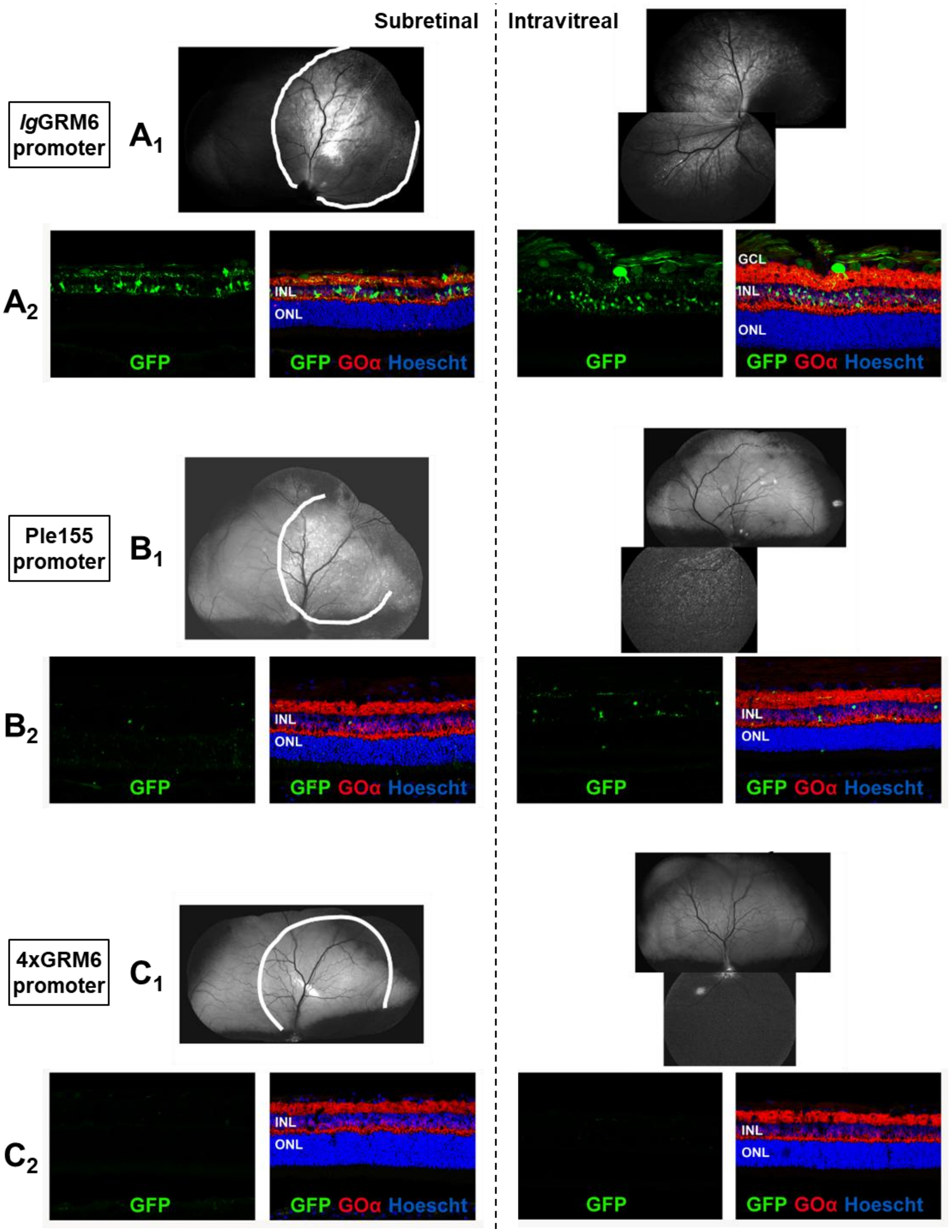


**Suppl. Fig. S2: Comparison of efficiency of three promoters at driving sfGFP expression in canine ON-BCs following subretinal and intravitreal injection of mixes of AAV capsid variants.**

**$A_1$, $B_1$, $C_1$**) Composite cSLO (BAF mode) images showing intense (*lg*GRM6 promoter, $A_1$), rare (Ple155 promoter, $B_1$), and lack of (4xGRM6 promoter, $C_1$) GFP fluorescence. White curved line shows border of subretinal bleb. **$A_2$, $B_2$, $C_2$**) Retinal cryosections showing native sfGFP fluorescence and Go$\alpha$ immunolabeling.

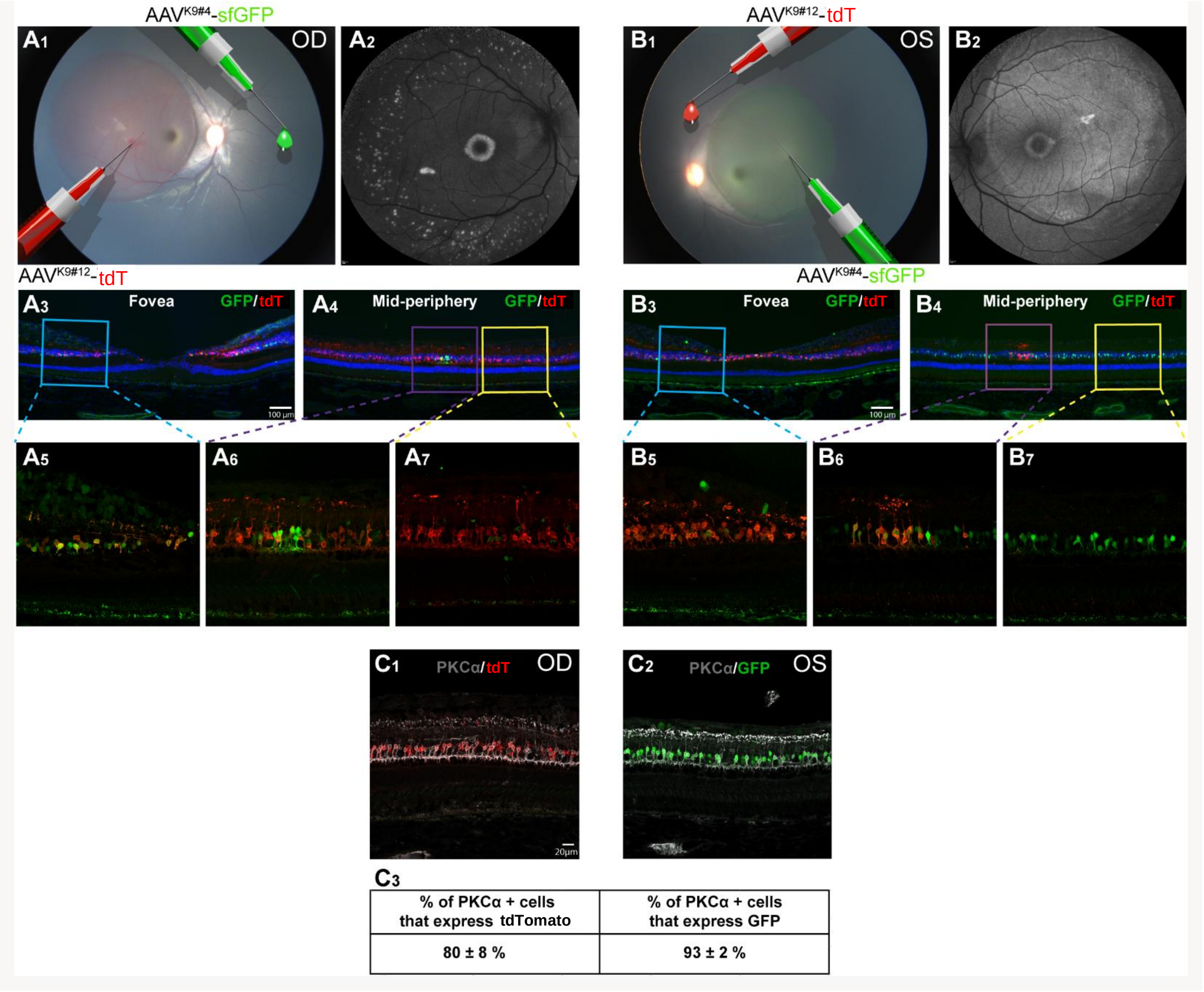


**Suppl. Fig. S3: Targeting of NHP ON-BCs with top performing AAV capsid variants.**

**$A_1$-$A_7$**) *In vivo* imaging and double fluorescence IHC of the right retina (OD) of a cynomolgus macaque injected intravitreally with AAV$^{K9\#4}$-*lg*GRM6-*sfGFP* and subretinally with AAV$^{K9\#12}$-*lg*GRM6-*tdTomato*. **$B_1$-$B_7$**) *In vivo* imaging and double fluorescence IHC of the left retina (OS) of the same animal injected intravitreally with AAV$^{K9\#12}$-*lg*GRM6-*tdTomato* and subretinally with AAV$^{K9\#4}$-*lg*GRM6-*sfGFP*. **$A_1$, $B_1$**) Modified fundus photograms of both retinas taken immediately after injections and pseudocolored to show routes of delivery (subretinal and intravitreal) of both viral vector constructs. **$A_2$, $B_2$**) cSLO images (BAF mode) showing detection of sfGFP fluorescence. **$A_3$, $B_3$**) Double IHC labeling of GFP and tdTomato at the fovea and perifovea. **$A_4$, $B_4$**) Double IHC labeling of GFP and tdTomato at the mid-periphery. **$C_1$**) Double IHC labeling of PKCα and tdTomato. **$C_2$**) Double IHC labeling of PKCα and GFP. **$C_3$**) Percentage (mean ± SD; n=4 replicates per eye) of PKCα-positive cells that co-expressed in the subretinally-treated (=bleb) area either tdTomato (in OD) or GFP (in OS).

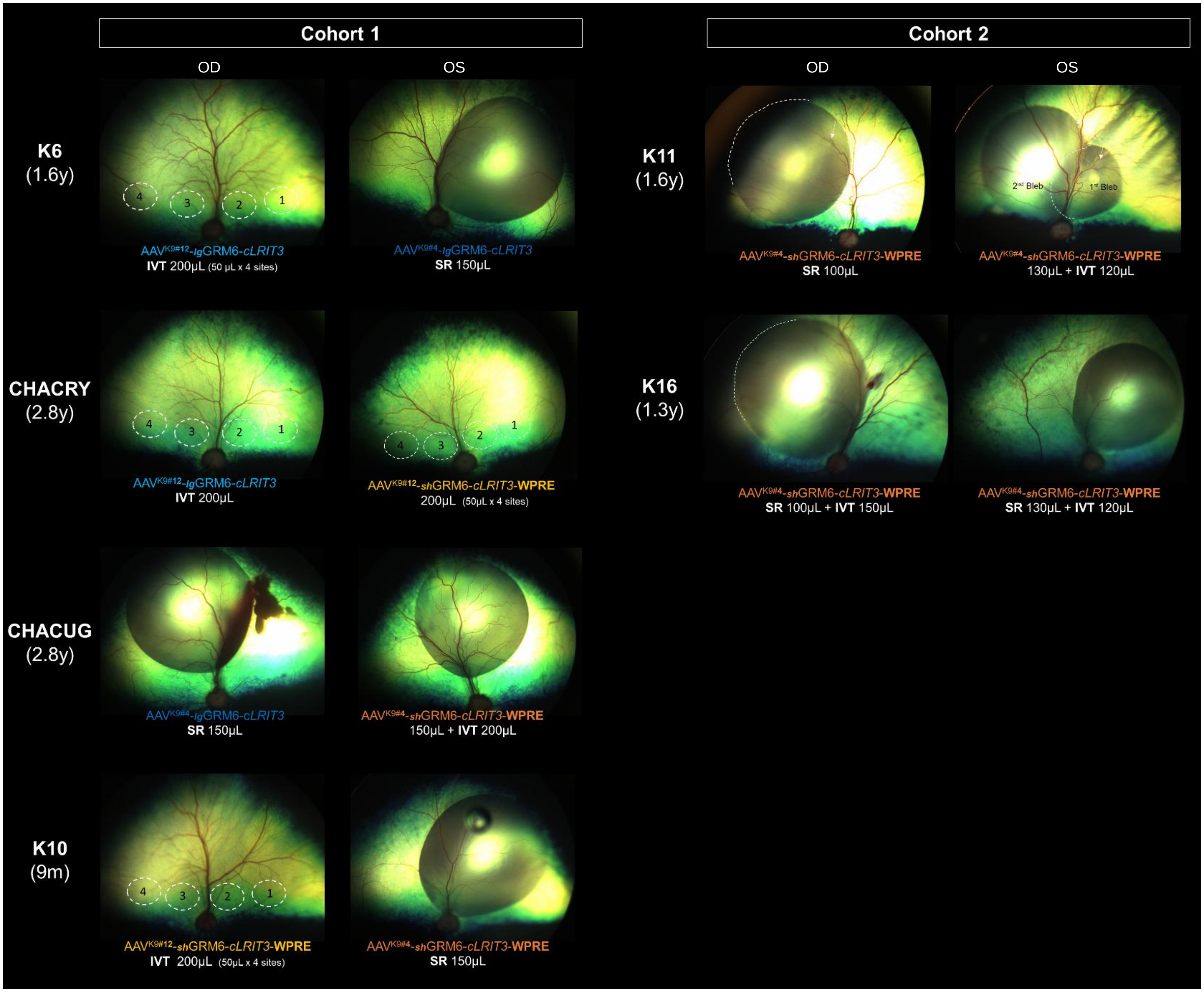


**Suppl. Fig. S4: Fundus pictures of dogs immediately after injection.**

*In vivo* imaging of *LRIT3*-CSNB dogs injected with AAV or BSS in each eye via subretinal (SR) or intravitreal (IVT) injection. SR injected eyes exhibit distinct subretinal blebs. The dotted white circles in IVT injected eyes indicate sites where doses were deposited pre-retinally. Hemorrhage in the right eye of CHACUG is due to inadvertent contact of subretinal injector needle with retinal vessels. OD, right eye; OS, left eye.

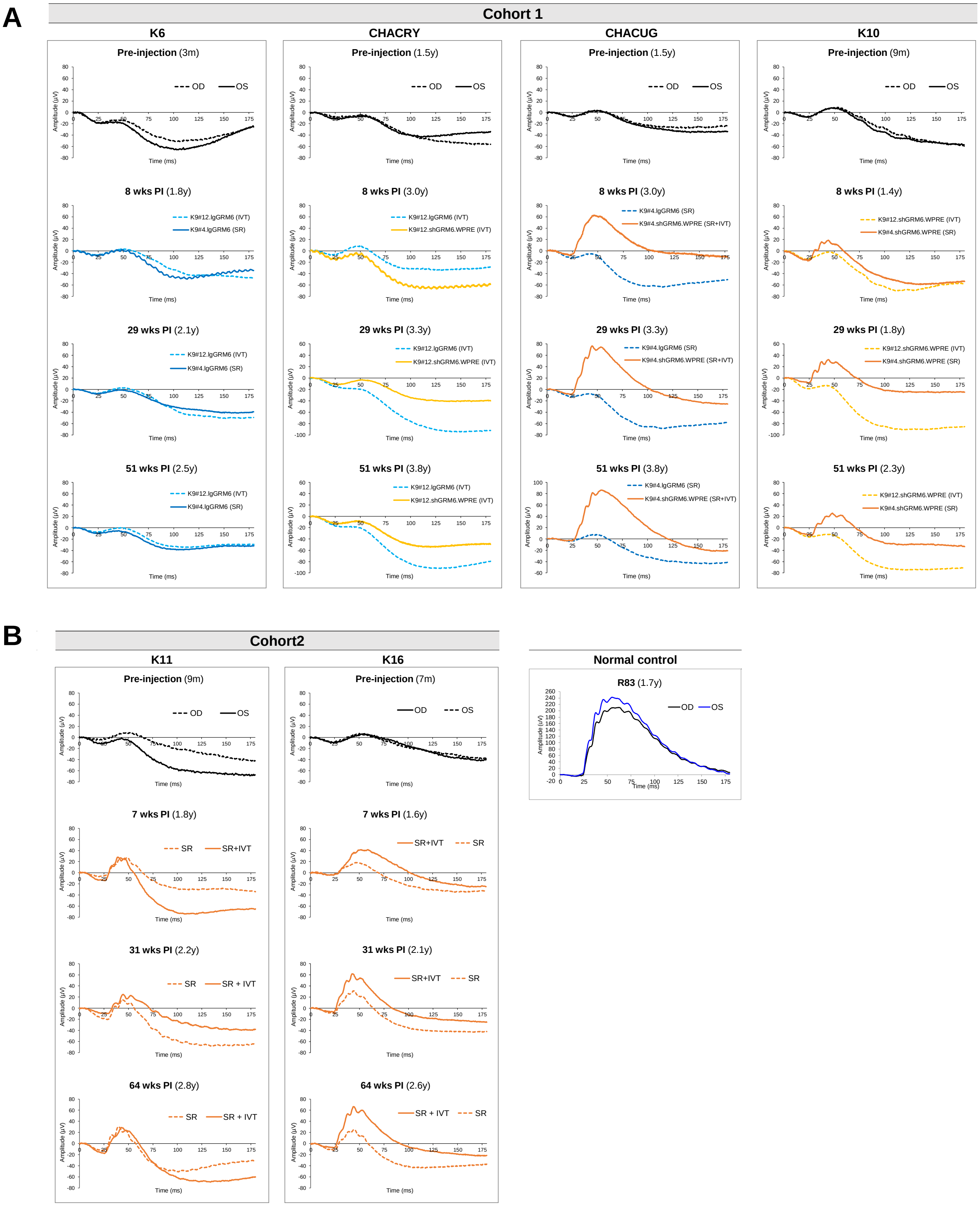


**Suppl. Fig. S5: Rod-derived ERG traces of individual dogs at selected time points.**

Rod-derived ERG traces of each eye in all dogs from Cohorts 1 (**A**) and 2 (**B**) at pre-injection, early (7-8 wks), mid (29-31 wks) and late (51-64 wks) post-injection time points. Post-injection traces are color-coded according to vectors injected, corresponding with the presentation in Table 1 and Suppl. Table S1. Age of dogs at ERG recording is shown in brackets. OD (dotted line), right eye; OS (solid line), left eye.

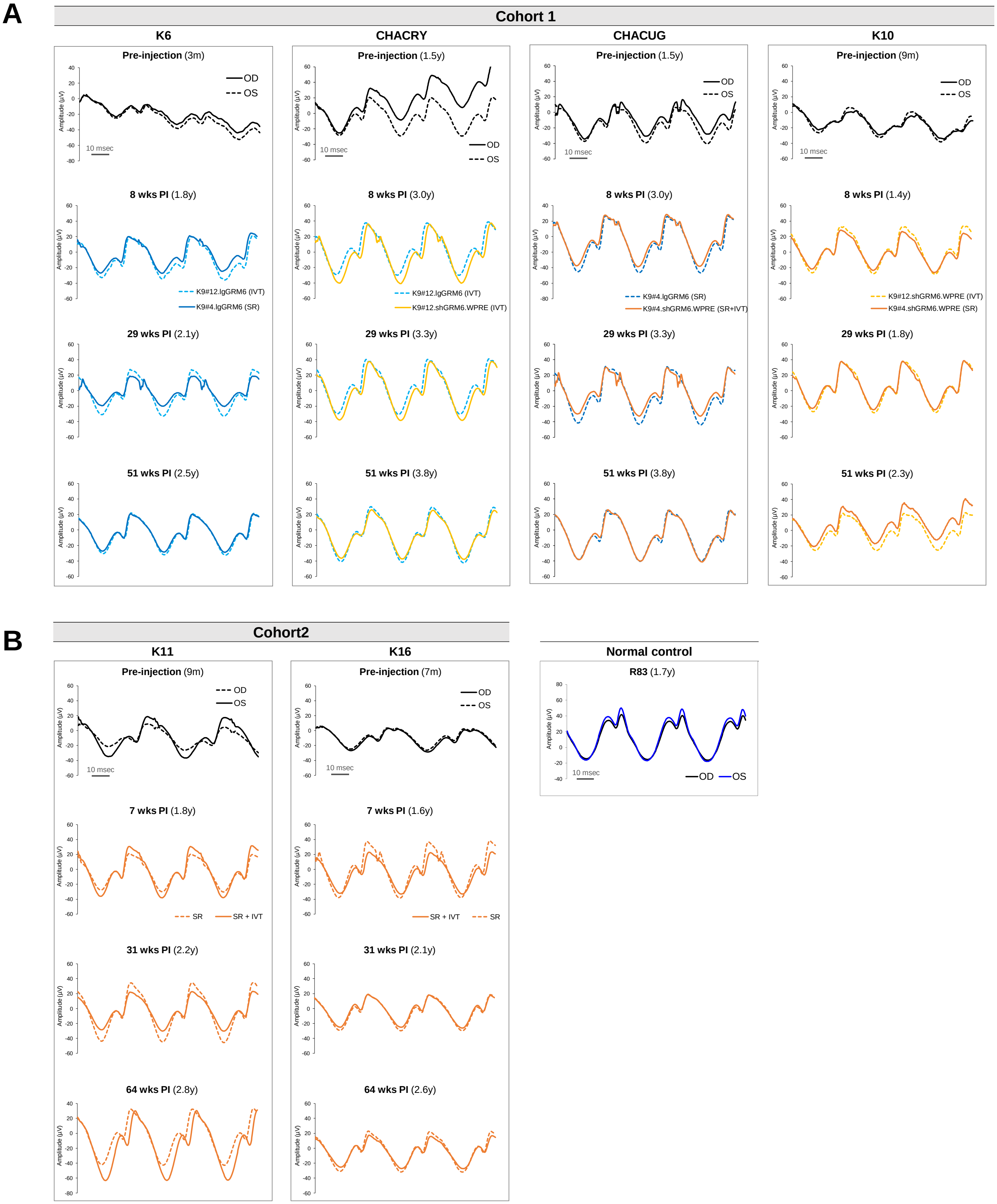


**Suppl. Fig. S6: Cone-derived ERG flicker traces of individual dogs at selected time points.** Cone-derived 29Hz ERG flicker traces of each eye in all the dogs from Cohorts 1 (**A**) and 2 (**B**) at pre-injection, early (7-8 wks), mid (29-31 wks) and late (51-64 wks) post-injection time points. The post-injection traces are color-coded according to the vectors injected, corresponding with the presentation in Table 1 and Suppl. Table S1. The age of dogs at ERG recording is shown in brackets. OD, right eye; OS, left eye.

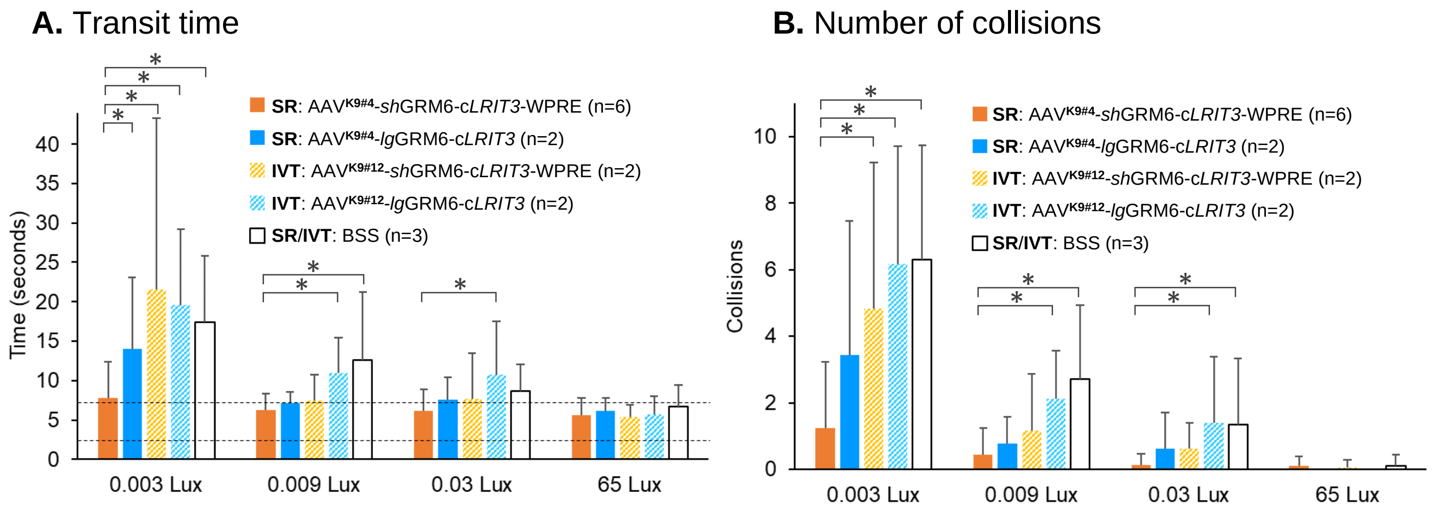


**Suppl. Fig. S7. Obstacle course performance per vector/route in injected CSNB dogs.**

Visually-guided navigation tested per eye in an obstacle course in six *LRIT3*-CSNB dogs (12 eyes) injected with one of the AAV vector variants in each eye. Visual function was assessed under dim scotopic (0.003, 0.09, 0.03 Lux) and ambient photopic (65 Lux) conditions. **A**) At the dimmest light intensity of 0.003 Lux, transit time was significantly reduced in eyes subretinally injected with the therapeutic vector (AAV$^{K9\#4}$-*sh*GRM6-c*LRIT3*-WPRE) (6 eyes) compared to eyes injected with either of the non-therapeutic vectors or BSS-injected control CSNB eyes (*p<.0001). Transition times at 65 Lux were minimally affected by the CSNB phenotype and, as such, comparable across the different therapeutic and control groups. The dotted lines define the 95% confidence interval of the transit time in 4 WT control dogs. **B**) Number of collisions at 0.003 Lux was significantly reduced in eyes subretinally injected with the therapeutic vector compared to eyes intravitreally injected with the non-therapeutic vectors, or BSS-injected control CSNB eyes (*p<.0001). While the bars represent the combined data of three available time points 18-50 weeks after the injection, there were no significant changes in vision testing outcomes over time. SR, subretinal; IVT, intravitreal.
